# Reduced expression of TCF7L2 in adipocyte impairs glucose tolerance associated with decreased insulin secretion, incretins levels and lipid metabolism dysregulation in male mice

**DOI:** 10.1101/2020.05.18.102384

**Authors:** Marie-Sophie Nguyen-Tu, Aida Martinez-Sanchez, Isabelle Leclerc, Guy A. Rutter, Gabriela da Silva Xavier

**Author notes:** Gabriela da Silva Xavier and Guy A. Rutter are joint corresponding authors. **Address:** Gabriela da Silva Xavier, Institute of Metabolism and Systems Research, IBR room 30, University of Birmingham, Edgbaston, Birmingham, B15 2TT, United Kingdom, Guy A. Rutter, Imperial College London, Section of Cell biology and Functional Genomics DuCane road, London W12 0NN, United Kingdom.

## Abstract

Transcription factor 7-like 2 (TCF7L2) is a downstream effector of the Wnt/beta-catenin signalling pathway and its expression is critical for adipocyte development. The precise role of TCF7L2 in glucose and lipid metabolism in adult adipocytes remains to be defined. Here, we aim to investigate how changes in TCF7L2 expression in mature adipocytes affect glucose homeostasis. *Tcf7l2* was selectively ablated from mature adipocytes in C57BL/6J mice using an adiponectin promoter-driven *Cre* recombinase to recombine alleles floxed at exon 1 of the *Tcf7l2* gene. Mice lacking *Tcf7l2* in mature adipocytes displayed normal body weight. Male mice exhibited normal glucose homeostasis at eight weeks of age. Male heterozygote knockout mice (aTCF7L2het) exhibited impaired glucose tolerance (AUC increased 1.14 ± 0.04 -fold, p=0.03), as assessed by intraperitoneal glucose tolerance test, and changes in fat mass at 16 weeks (increased by 1.4 ± 0.09-fold, p=0.007). Homozygote knockout mice exhibited impaired oral glucose tolerance at 16 weeks of age (AUC increased 2.15 ± 0.15-fold, p=0.0001). Islets of Langerhans exhibited impaired glucose-stimulated insulin secretion *in vitro* (decreased 0.54 ± 0.13-fold aTCF7L2KO vs control, p=0.02), but no changes in *in vivo* glucose-stimulated insulin secretion. Female mice in which one or two alleles of the *Tcf7l2* gene was knocked out in adipocytes displayed no changes in glucose tolerance, insulin sensitivity or insulin secretion. Plasma levels of glucagon-like peptide-1 and gastric inhibitory polypeptide were lowered in knockout mice (decreased 0.57 ± 0.03-fold and 0.41 ± 0.12-fold, p=0.04 and p=0.002, respectively), whilst plasma free fatty acids and Fatty Acid Binding Protein 4 circulating levels were increased by 1.27 ± 0.07 and 1.78 ± 0.32-fold, respectively (p=0.05 and p=0.03). Mice with biallelic *Tcf7l2* deletion exposed to high fat diet for 9 weeks exhibited impaired glucose tolerance (p=0.003 at 15 min after glucose injection) which was associated with reduced *in vivo* glucose-stimulated insulin secretion (decreased 0.51 ± 0.03-fold, p=0.02). Thus, our data indicate that loss of *Tcf7l2* gene expression in adipocytes leads to impairments on metabolic responses which are dependent on gender, age and nutritional status. Our findings further illuminate the role of TCF7L2 in the maintenance of glucose homeostasis.

## Introduction

Transcription factor 7-like 2 (TCF7L2) is a member of the high mobility group box family of transcription factors, a downstream effector of the Wnt/beta-catenin signalling pathway, and a key regulator of development and cell growth [1]. TCF7L2 function is important for the proper function of tissues involved in the regulation of energy homeostasis. For example, TCF7L2 is required for the maintenance of functional pancreatic beta cell mass and insulin release from the endocrine pancreas [2, 3]. In the liver, ablation of *Tcf7l2* expression from hepatocytes has variously been shown to lead to reduced hepatic glucose production and improved glucose homeostasis [4], or hyperglycemia [5]. Studies by MacDougald and colleagues [6–8] have demonstrated that *Tcf7l2* is involved in the regulation of the expression of proadipogenic genes during adipocyte development. Additionally, insulin/insulin receptor substrate-1 and insulin growth factor 1 have been shown to cross-talk with the Wnt signalling pathway to regulate insulin sensitivity in preadipocytes [9, 10]. The presence of TCF7L2 binding sites on the promoter of the insulin receptor gene also suggests a role of TCF7L2/Wnt signalling pathway in regulating insulin action in adipocytes [11].

Genome-wide association studies (GWAS) have identified single nucleotide polymorphisms (SNPs; rs12255372 and rs7903146) in the *TCF7L2* gene as being amongst the most strongly associated with an increased risk of type 2 diabetes [12]. Humans carrying the T risk allele at rs7903146 have elevated proinsulin levels, lowered first-phase insulin secretion and impaired responses to the incretin hormone glucagon-like peptide 1 (GLP-1) [13–17]. Whilst *TCF7L2* variants appear chiefly to affect pancreatic beta cell function in man, the mechanisms driving impaired insulin secretion are still poorly defined [18, 19]. Although rs7903146 in *TCF7L2* is not associated with changes in overall *TCF7L2* transcription (i.e. of all isoforms), with conflicting data regarding the association of rs7903146 with specific *TCF7L2* variant expression in subcutaneous fat [20–22], *TCF7L2* expression has been shown to be reduced in adipose tissue from type 2 diabetes subjects [23] and in obese mice [24]. Additionally, surgery-induced weight loss has been shown to regulate alternative splicing of *Tcf7l2* in adipose tissue [25]. TCF7L2 splice variant expression is regulated by plasma triglycerides and free fatty acids [25, 26], with evidence indicating that acute intake of fat leads to reduced expression of *TCF7L2* in human adipocytes [27]. Thus, overall these data suggest that changes in *TCF7L2* expression may be linked to adaptation to changes in fuel intake.

Efforts to understand the causal relationship between TCF7L2 and diabetes led to studies on several metabolic tissues, with the data suggesting that the combined effects of loss of TCF7L2 in multiple tissues may account for the diabetes phenotype [28]. For example, previous reports have described the impact of loss of *TCF7L2* expression on adipocyte development and insulin sensitivity [24, 29]. In the present study, we used a genetic model of ablation of *TCF7L2* gene expression in mature adipocytes to determine whether TCF7L2 plays a role in adipocyte function, independent of its role in adipogenesis. We have focused on whether loss of *TCF7L2* expression in the adipocyte impacts the release of hormones involved in glucose homeostasis, notably insulin and incretins. In this way, we sought to explore the possibility that altered *TCF7L2* expression in the adipocyte may contribute to type 2 diabetes risk through downstream effects on multiple effector organs [28, 30].

## Methods

### Animal care and maintenance

To generate tissue-specific knockout of *Tcf7l2* alleles, we crossed mice in which exon 1 (encoding for the beta-catenin-binding domain) was flanked by *LoxP* sites [2] with mice expressing *Cre* recombinase under the control of the adiponectin promoter [31] (a kind gift from D. Withers, Imperial College London) to produce deletion of one *Tcf7l2* allele (aTCF7L2het) or two alleles (aTCF7L2KO). Littermates used as controls did not express *Cre* recombinase but were homozygous or heterozygous for the floxed *Tcf7l2* allele. Adiponectin-*Cre* [31] or mice with *Tcf7l2* gene flanked by LoxP sites (TCF7L2-floxed) [2] did not display phenotypes that deviate from wild-type littermate control mice, consequently we used TCF7L2-floxed mice as controls in our test cohorts. Mice were born at the expected Mendelian ratios with no apparent abnormalities. Animals were housed 2-5 per individually-ventilated cage in a pathogen-free facility with 12:12 light:dark cycle with free access to standard mouse chow (RM-1; Special Diet Services) diet and water. High fat diet (HFD) cohort were put under a high sucrose high fat diet (D12331; Research diets) for 12 weeks from 7-week-old. For the chow diet cohort, metabolic exploration was performed on each animal within a 2-week window at 2 stages (8-week-old and 16-week-old). All *in vivo* procedures described were performed at the Imperial College Central Biomedical Service and approved by the UK Home Office Animals Scientific Procedures Act, 1986 (HO Licence PPL 70/7971 to GdSX).

### Metabolic tolerance tests

Glucose tolerance was performed on 15 h-fasted mice after an oral gavage of glucose (OGTT, 2 g/kg of body weight) or intraperitoneal injection of glucose (IPGTT, 1g/kg body weight. IPGTT and OGTT were performed at two stages (at 8-week-old and at 16-week-old) for each individual mouse. Insulin tolerance was performed after a 5 h-fast with an intraperitoneal injection of insulin (IPITT, 0.5 U/kg in females, 0.75 U/kg in males under chow diet, 1.5U/kg in males under HFD). *In vivo* glucose-stimulated insulin secretion was assessed after oral or intraperitoneal administration of glucose and blood was collected at 0- and 15-minutes post-injection to assess plasma insulin levels using an ultra-sensitive mouse insulin ELISA kit (Crystal Chem, Netherlands) or using a Homogeneous Time Resolved Fluorescence (HTRF) insulin kit (Cisbio, France) in a PHERAstar reader (BMG Labtech, UK).

### Protein isolation and Western immuno-blotting

To assess insulin sensitivity, male mice were starved for four hours and following an intraperitoneal injection of insulin (1UI/kg body weight) adipose tissues were collected and frozen in liquid nitrogen. Adipose tissue proteins were extracted in lysis buffer (150 mmol/l NaCl, 50 mmol/l Tris-HCl pH 8.0, 1% NP-40) supplemented with protease inhibitors (Roche, Germany) and phosphatase inhibitors (Sigma-Aldrich, UK) and analysed by Western blotting using antibodies for TCF4/TCF7L2 (C48H11) (#2569, 1:500, Cell signalling, NEB, UK), phospho-AKT (#9271, 1:1000, Cell signalling, NEB, UK), total-AKT (#9272, 1:1000, Cell signalling, NEB, UK), GAPDH (#2118, 1:10000, Cell signalling, NEB, UK), alpha-tubulin (T5168, 1:10 000, Sigma-Aldrich, UK). Fiji software was used for densitometry quantification. Uncut versions of all Western blot images are presented in supplemental figure S1.

### Primary islet studies

Islets were isolated by digestion with collagenase as described [32]. In brief, pancreata were inflated with a solution of collagenase from *clostridium histolyticum* (1 mg/mL; Nordmark, Germany) and placed in a water bath at 37 ⁰C for 12 min. Islets were washed and purified on a Histopaque gradient (Sigma-Aldrich, UK). Isolated islets were cultured for 24 h in RPMI 1640 containing 11.1 mmol/l glucose, 10% foetal bovine serum and L-glutamine (Sigma-Aldrich, UK) and allowed to recover overnight.

Insulin secretion assays on isolated mouse islets were performed as previously described [32]. In brief, 10 size-matched islets per condition were incubated for 1h in Krebs-HEPES-bicarbonate (KHB) solution (130 mmol/L NaCl, 3.6 mmol/L KCl, 1.5 mmol/L CaCl_2_, 0.5 mmol/L MgSO_4_, 0.5 mmol/L KH_2_PO_4_, 2 mmol/L NaHCO_3_, 10 mmol/L HEPES, and 0.1% BSA, pH 7.4) containing 3 mmol/L glucose. Subsequently, islets were incubated for 30 minutes in KHB solution with either 3 mmol/L-glucose, 17 mmol/L-glucose or 30 mmol/L-KCl. Secreted and total insulin were quantified using a Homogeneous Time Resolved Fluorescence (HTRF) insulin kit (Cisbio, France) in a PHERAstar reader (BMG Labtech, UK) following the manufacturer’s guidelines.

Measurement of intracellular calcium dynamics was performed as previously described [33]. In brief, whole isolated islets were incubated with fura-8AM (Invitrogen, UK) [34] for 45 min at 37⁰C in KHB containing 3 mmol/L glucose. Fluorescence imaging was performed using a Nipkow spinning disk head, allowing rapid scanning of islet areas for prolonged periods of time with minimal phototoxicity. Volocity software (PerkinElmer Life Sciences, UK) provided interface while islets were kept at 37⁰C and constantly perifused with KHB containing 3 mmol/L or 17 mmol/L glucose or 30 mmol/L KCl.

### Histology

Epididymal adipose tissue were removed from euthanized mice, fixed overnight in 10% formalin and subsequently embedded in paraffin wax. Adipose tissue slices (5 μm) were stained with hematoxylin and eosin (Sigma-Aldrich, UK) for morphological analysis.

### Analysis of circulating factors in plasma and serum

Blood in the fed state was obtained by tail bleeding, and circulating factor concentrations were measured using the following kits according to the respective manufacturer’s protocols: Bio-Plex protein array system (Biorad, UK) with one multiplex panel was used to measure total Glucagon-like peptide-1 (GLP-1), glucose-dependent insulinotropic polypeptide (GIP), leptin, adiponectin and plasminogen activator inhibitor-1 (PAI-1); one multiplex panel was used to measure fatty acid binding protein 4 (FABP4) and resistin (R&D Systems, UK); non-esterified fatty acid (NEFA) serum levels were measured by colorimetric assay (Randox, UK) and dipeptidyl peptidase 4 (DPP4; R&D Systems, UK) levels were measured by ELISA.

### RNA isolation and quantitative PCR

RNA was isolated from epididymal and subcutaneous adipose tissue, liver and pancreatic islets with TRIzol following manufacturer’s instructions (Invitrogen, UK). RNA purity and concentration were measured by spectrophotometry (Nanodrop, Thermo Scientific, UK). Only RNA with absorption ratios between 1.8-2.0 for 260/280 and 260/230nm were used. RNA integrity was checked on an agarose gel. RNA was reversed transcribed using High-Capacity cDNA Reverse Transcription kit (Applied Biosystems, UK). qPCR was performed with Fast SYBR green master mix (Applied Biosystems, UK). The comparative Ct method (2^−ΔΔCT^) was used to calculate relative gene expression levels using *Gapdh*, *βactin* or *Ppia* as an internal control. The primers sequences are listed in Supplemental table S1.

### Statistical analysis

Data are shown as means ± SEM. GraphPad Prism 8.4 was used for statistical analysis. Statistical significance was evaluated by the two-tailed unpaired Student t-test and one- or two-way ANOVA, with Tukey or Bonferroni multiple comparisons post-hoc test as indicated in the figure legends. P values of <0.05 were considered statistically significant.

## Results

### Effects of adipocyte-selective Tcf7l2 deletion on body weight and fat mass

To determine the role of TCF7L2 in the mature adipocyte, we generated a mouse line in which *Tcf7l2* is deleted specifically in these cells through the expression of *Cre* recombinase under the control of the adiponectin promoter [31]. *Cre* recombinase mediated the excision of exon 1 of *Tcf7l2* generating one single *Tcf7l2* allele deletion (aTCF7L2het) or a biallelic *Tcf7l2* deletion (aTCF7L2KO). *Tcf7l2* mRNA levels were decreased in inguinal adipose tissue (iWAT) by 39.8 ± 13.3% (p=0.008) while in epididymal adipose tissue (eWAT), a reduction by 46.3 ± 20.1% did not reach any statistical significance (p=0.09) in aTCF7L2het mice compared to controls. In aTCF7L2KO mice, expression was reduced by 77.3 ± 11.2 % and 57.8 ± 20.2%, respectively in eWAT and iWAT compared to controls. Conversely, no changes in *Tcf7l2* expression were apparent in liver and pancreatic islets from aTCF7L2het and aTCF7L2KO mice vs islets from littermate controls (Fig.1a). Correspondingly, the content of the two TCF7L2 protein isoforms (79kDa and 58kDa) in eWAT was significantly reduced by 41 ± 11 % and 35 ± 7 % respectively in aTCF7L2het mice and reduced by 77 ± 17 % and 80 ± 14 % in aTCF7L2KO mice (Fig.1b and c). Body weight was unchanged in male (Fig.1d) and female (Fig.1e) in aTCF7L2het and aTCF7L2KO mice maintained on normal chow diet (NC). No apparent change in adipocyte morphology was observed in male mice maintained under NC (Fig.1f). Likewise, we found no change in fat or lean mass in 8 weeks old male or female mice, as assessed by echoMRI (Fig.1g, i, k and m). However, fat mass was significantly increased, and lean mass was decreased, in male aTCF7L2het mice on normal chow diet (Fig.1h and j). Female mice showed no change in body composition with age (Fig.1l and n).

**Fig. 1.**
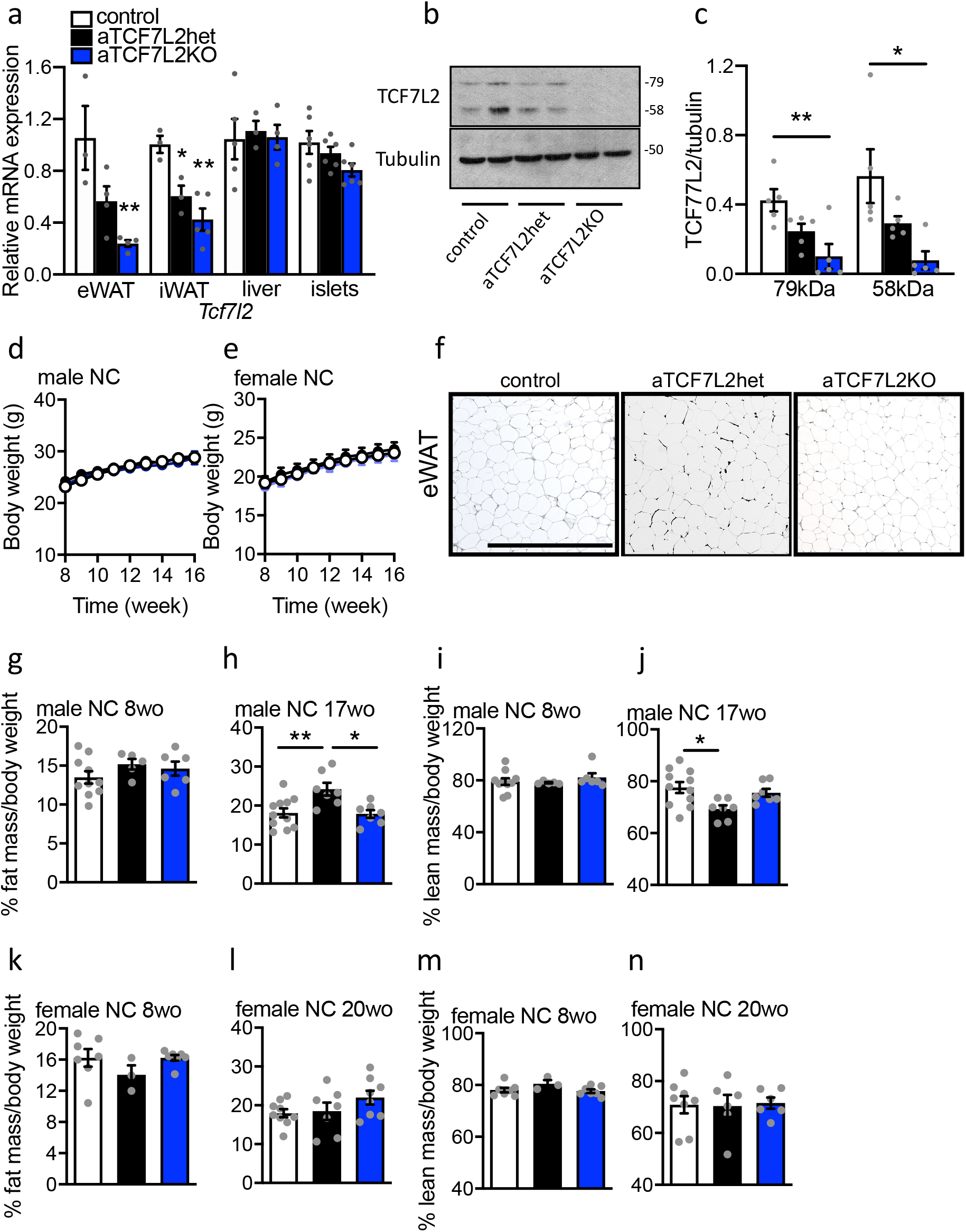
Effects of adipocyte-selective *Tcf7l2* deletion on body weight and fat mass in males and females under normal chow diet. (**a**) *Tcf7l2* mRNA expression by RT-qPCR in epididymal white adipose tissue (eWAT), inguinal subcutaneous adipose tissue (iWAT), liver and isolated islets (n=3-6 mice/genotype). One-way ANOVA followed by Tukey post hoc test **p<0.01 and *p<0.05 versus control. (**b**) TCF7L2 protein expression in eWAT by western blotting. (**c**) Densitometry quantification TCF7L2 expression by western blotting in eWAT (n= 4 mice/genotype). One-way ANOVA followed by Tukey post hoc test **p<0.01 and *p<0.05 versus control. (**d**) Body weight in males under chow diet (NC) (n=10 control mice, n=9 aTCF7L2het mice, n=6 aTCF7L2KO mice). (**e**) Body weight in females under NC (n=13 control mice, n=7 aTCF7L2het mice, n=7 aTCF7L2KO mice). (**f**) Hematoxylin and Eosin Staining of eWAT in normal chow diet NC. Scale bar is 100 μm. (**g**) Percentage fat mass over body weight in 8-week-old males (n=10 control mice, n=5 aTCF7L2het mice, n=6 aTCF7L2KO mice). (**h**) Percentage fat mass over body weight in 17-week-old males (n=11 control mice, n=6 aTCF7L2het mice, n=7 aTCF7L2KO mice). One-way ANOVA followed by Tukey post hoc test **p<0.01 versus control, *p<0.05 versus aTCF7L2het. (**i**) Percentage lean mass over body weight in 8-week-old males (n=10 control mice, n=5 aTCF7L2het mice, n=6 aTCF7L2KO mice). (**j**) Percentage lean mass over body weight in 17-week-old males (n=11 control mice, n=6 aTCF7L2het mice, n=7 aTCF7L2KO mice). One-way ANOVA followed by Tukey post hoc test *p<0.05 versus control. (**k**) Percentage fat mass over body weight in 8-week-old females (n=7 control mice, n=3 aTCF7L2het mice, n=7 aTCF7L2KO mice). (**l**) Percentage fat mass over body weight in 17-week-old females (n=9 control mice, n=7 aTCF7L2het mice, n=8 aTCF7L2KO mice). (**m**) Percentage lean mass over body weight male in 8-week-old females (n=7 control mice, n=3 aTCF7L2het mice, n=7 aTCF7L2KO mice). (**n**) Percentage lean mass over body weight in 17-week-old females (n=9 control mice, n=7 aTCF7L2het mice, n=8 aTCF7L2KO mice). Data shown as mean ± SEM.

### Effects of adipocyte-selective Tcf7l2 deletion on glucose tolerance and insulin sensitivity

To explore the effects of adipocyte-specific *Tcf7l2* ablation on whole body metabolism, we measured glucose tolerance, and found an age-dependent impairment in the response to glucose administration. Glucose challenge was assessed in male and female mice at a young stage (8 weeks of age) and at an older stage (16 weeks of age). In younger mice, blood glucose levels after intraperitoneal injection of glucose were similar in aTCF7L2het and aTCF7L2KO compared to sex-matched littermate controls regardless of the gender (Fig.2a and Fig.3a). In older mice, glucose levels were impaired in male aTCF7L2het mice rising to 17.7 ± 1.0 mmol/L at 15 minutes after intraperitoneal injection of glucose compared to littermate controls rising to 13.4 ± 0.8 mmol/L (Fig.2b). Likewise, aTCF7L2het and young aTCF7L2KO mice had similar increases in glucose levels 15 minutes after oral administration of glucose (17.4 ± 1.1 mmol/L and 15.8 ± 0.7 mmol/L respectively) compared to littermate controls (16.7 ± 0.7 mmol/L; Fig.2c). Surprisingly, glucose tolerance (as assessed by oral glucose tolerance test) was impaired in older aTCF7L2KO mice (17 weeks) compared to littermate controls (Fig.2d). Female mice exhibited no change in glucose tolerance regardless of age and genotype (Fig.3a, Fig.3b and Fig.3c). Whole body insulin sensitivity was unaffected in both genders across all genotypes (Fig.2e and Fig.3d).

**Fig. 2.**
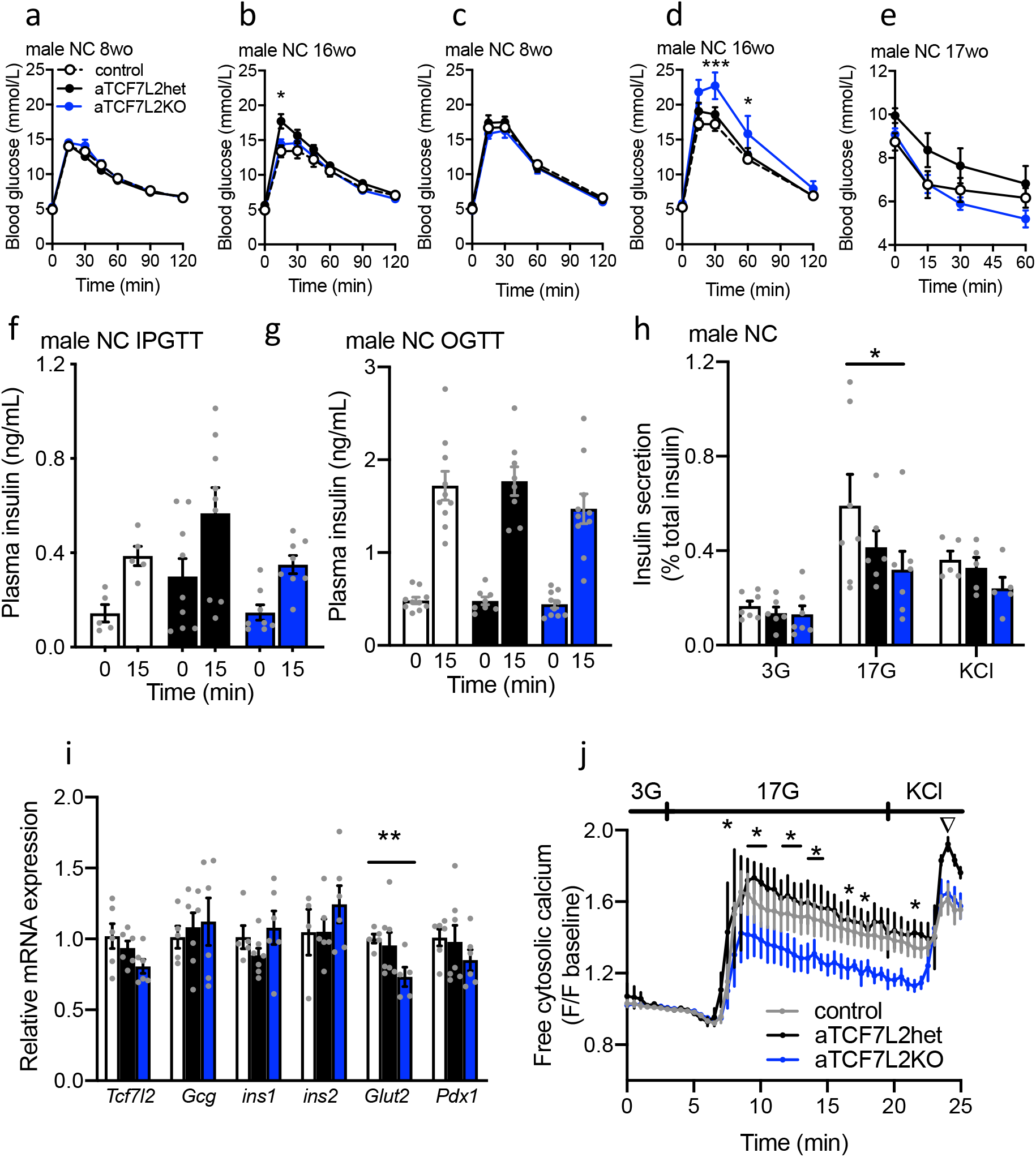
Effects of adipocyte-selective *Tcf7l2* deletion on glucose tolerance and beta cell function in male mice maintained on a normal chow diet. (**a**) Blood glucose levels during an intraperitoneal glucose tolerance test (IPGTT) in 8-week-old male mice (n=10 control mice, n=10 aTCF7L2het mice, n=9 aTCF7L2KO mice) maintained on a normal chow diet (NC). (**b**) Blood glucose levels during IPGTT in 16-week-old male mice (n=10 control mice, n=10 aTCF7L2het mice, n=9 aTCF7L2KO mice). Data were analysed by two-way ANOVA followed by Bonferroni post-hoc test, *p<0.05 aTCF7L2het versus control group. Blood glucose levels during oral glucose tolerance test (OGTT) in 8-week-old male mice (**c**; n=10 control mice, n=6 aTCF7L2het mice, n=6 aTCF7L2KO mice), and 16-week-old males (**d**; n=10 control mice, n=6 aTCF7L2het mice, n=6 aTCF7L2KO mice). Data were analysed by two-way ANOVA followed by Bonferroni post-hoc test ***p<0.001 and *p<0.05 aTCF7L2KO versus control. Blood glucose levels during IPITT in 17-week-old male mice (**e;**n=10 control mice, n=10 aTCF7L2het mice, n=10 aTCF7L2KO mice). Plasma insulin levels **(f;**n=5 control mice, n=9 aTCF7L2het mice, n=8 aTCF7L2KO mice) during IPGTT after intraperitoneal injection of glucose (3g/kg) in 17-week-old male mice. Data in (e) were analysed by two-way ANOVA followed by Bonferroni post-hoc test with no statistical significance between parameters (**g**) Insulin plasma levels during OGTT (3g/kg) in 17-week-old male mice (n=10 control mice, n=7 aTCF7L2het mice, n=9 aTCF7L2KO mice). Data were analysed with Student’s t-test, with no statistical significance between parameters. (**h**) Insulin secretion was measured by static incubation on isolated islets from 17-week-old male mice. Data were analysed by two-way ANOVA followed by Tukey post-hoc test *p<0.05 aTCF7L2KO versus control, n=5-7 mice/genotype. (**i**) mRNA expression profiling by RT-qPCR of key pancreatic islet markers in isolated islets from 17-week-old male mice; each dot represents data from one mouse. Data were analysed using Student’s t-test, **p<0.01 aTCF7L2KO versus control. (**j**) Measurement of dynamic changes in intracellular calcium concentrations in isolated islets from 17-week-old male mice in response to perfusion of low (3 mmol/l, 3G), high (17 mmol/l, 17G) glucose and 30 mmol/l KCl and represented as fold change of fluorescence intensity compared to basal state at low glucose (n=3 mice/genotype). Two-way ANOVA followed by Tukey post-hoc test, *p<0.05 aTCF7L2het versus aTCF7L2KO, ▽p<0.05 aTCF7L2het versus control. Data are shown as mean ± SEM.

**Fig. 3.**
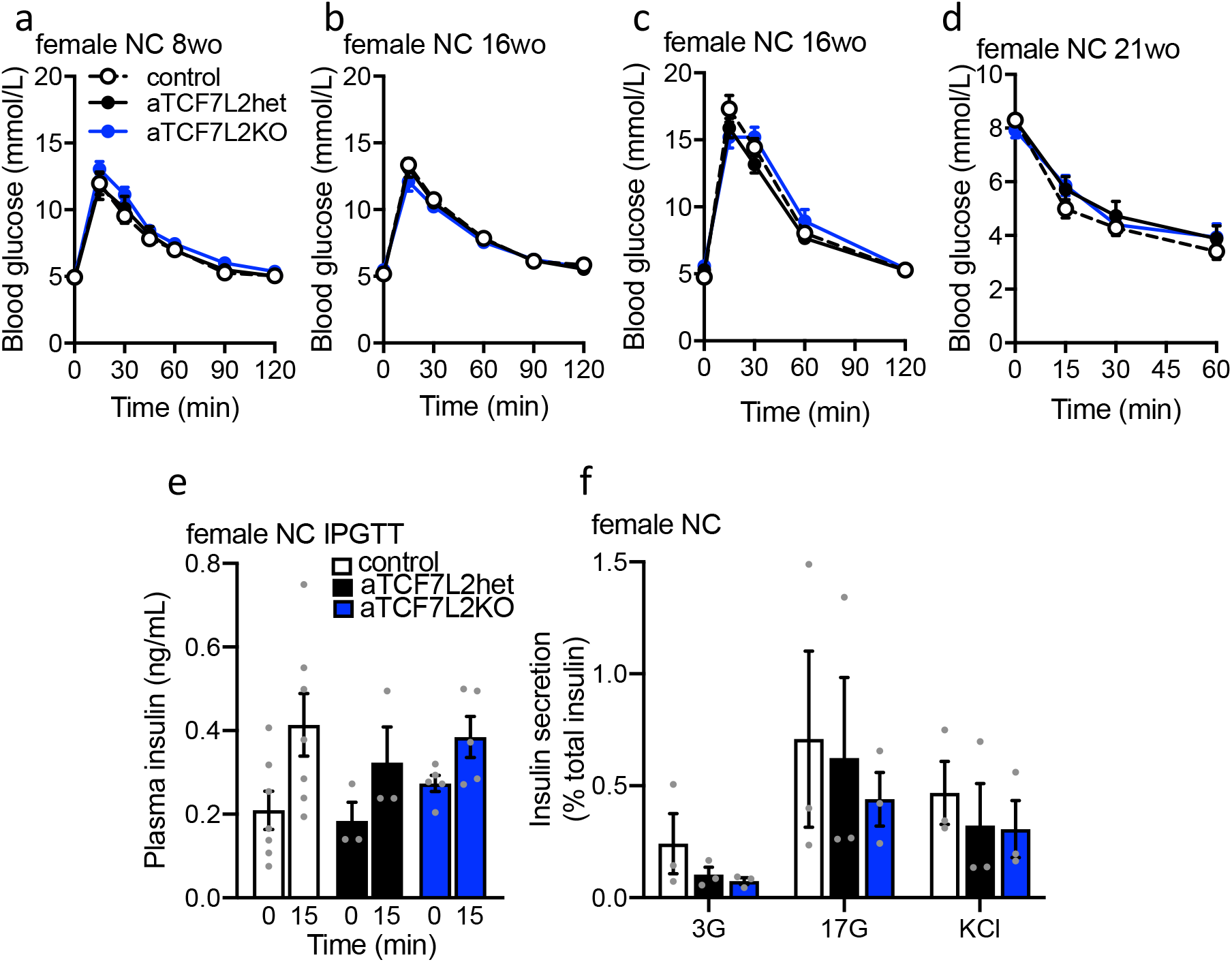
Effects of adipocyte-selective *Tcf7l2* deletion on glucose tolerance and beta cell function in female mice maintained on a normal chow diet. (**a**) Blood glucose levels during IPGTT in 8-week-old female mice (n=10 control mice, n=10 aTCF7L2het mice, n=8 aTCF7L2KO mice) maintained on a normal chow diet (NC). (**b**) Blood glucose levels during IPGTT in 16-week-old female mice (n=10 control mice, n=10 aTCF7L2het mice, n=8 aTCF7L2KO mice). (**c**) Blood glucose levels during OGTT in 16-week-old female mice (n=6 control mice, n=9 aTCF7L2het mice, n=4 aTCF7L2KO mice). (**d**) Blood glucose levels during IPITT in 21-week-old female mice (n=10 control mice, n=7 aTCF7L2het mice, n=5 aTCF7L2KO mice). **(e**) Insulin plasma levels during IPGTT (3g/kg) in 20-week-old female mice under chow diet (n=7 control mice, n=3 aTCF7L2het mice, n=5 aTCF7L2KO mice). (**f**) Insulin secretion on isolated islets from 20-week-old female mice during static incubation of 3 mmol/l glucose (3G), 17 mmol/l glucose (17G) and 30 mmol/l KCl, (n=3 mice/genotype). Data shown as mean ± SEM.

### Effects of adipocyte-selective Tcf7l2 deletion on pancreatic beta cell function

In order to assess whether ablation of TCF7L2 in mature adipocyte affects organs involved in glycaemic control, beta cell secretory capacity was measured during glucose challenge. We sought first to examine whether impaired intraperitoneal and oral glucose challenge in older male aTCF7L2het and aTCF7L2KO mice was due to an impact on insulin secretion. Fasting plasma insulin levels and *in vivo* glucose-stimulated insulin release were similar in aTCF7L2het and aTCF7L2KO mice vs littermate controls after intraperitoneal injection of glucose (Fig.2f) or after oral administration of glucose (Fig.2g). Glucose stimulated insulin secretion in isolated islets was decreased after high (17 mM) glucose incubation (0.32 ± 0.08% in aTCF7L2KO vs 0.59 ± 0.13% in controls) while responses to KCl (30 mM) were not different in islets from aTCF7L2KO compared to islets from littermate controls (Fig.2h).

When exploring after TCF7L2 loss expression of key genes associated with normal beta cell function, we observed no significant changes in gene expression. Indeed, no difference was observed for the expression of the insulin (*Ins1, Ins2*), or glucagon (*Gcg*) genes in islets from all genotypes, however a reduction in the expression of the glucose transporter 2 (Glut2/*Slc2a2*) gene was observed in aTCF7L2KO mice (0.73 ± 0.06 in aTCF7L2KO vs 1.00 ± 0.03 in controls; Fig.2i). To further evaluate the origins of the secretory defects observed in isolated islets (Fig. 2h), we measured the changes in cytosolic calcium in response to incubation with varying concentrations of glucose (3 mM, 17 mM) in the presence or absence of KCl (30 mM) on isolated islets (Fig.2j). Islets from aTCF7L2KO male mice showed a diminished response to high glucose incubation compared to aTCF7L2het animals, while differences compared to control mice did not reach any statistical significance. Islets from aTCF7L2het male mice showed an increase response to KCl compared to controls (Fig.2j). *In vivo* after intraperitoneal injection and *in vitro* glucose stimulated insulin secretion was unchanged in female aTCF7L2KOhet and aTCF7L2KO mice (Fig.3e and f). Therefore, alterations in pancreatic beta cell function observed *ex vivo* in the absence of TCF7L2 in adipocyte have no impact on whole body glucose-stimulated insulin secretion.

### Plasma levels of incretins and free fatty acids depend on adipocyte TCF7L2 expression

To investigate the causes of impaired oral glucose tolerance in male aTCF7L2KO mice, we measured the circulating levels of other hormones regulating glucose metabolism. Circulating GLP-1 and GIP levels in plasma were decreased in older male aTCF7L2KO mice compared to age-and sex-matched littermate control mice (GLP-1: 23.7 ± 6.8 vs 57.6 ± 11.6 ng/mL; GIP: 335.1 ± 21.4 vs 583.8 ± 39.5 ng/mL respectively; Fig.4a and Fig.4b). We observed a significant decrease in GIP levels (421.8 ± 24.6 vs 583.8 ± 39.5 ng/mL in controls), but no robust or statistical differences in GLP-1 levels in aTCF7L2het mice (Fig.4a and b). Plasma DPP4 levels in male aTCF7L2KO mice were not different compared to controls (Fig.4c).

**Fig. 4.**
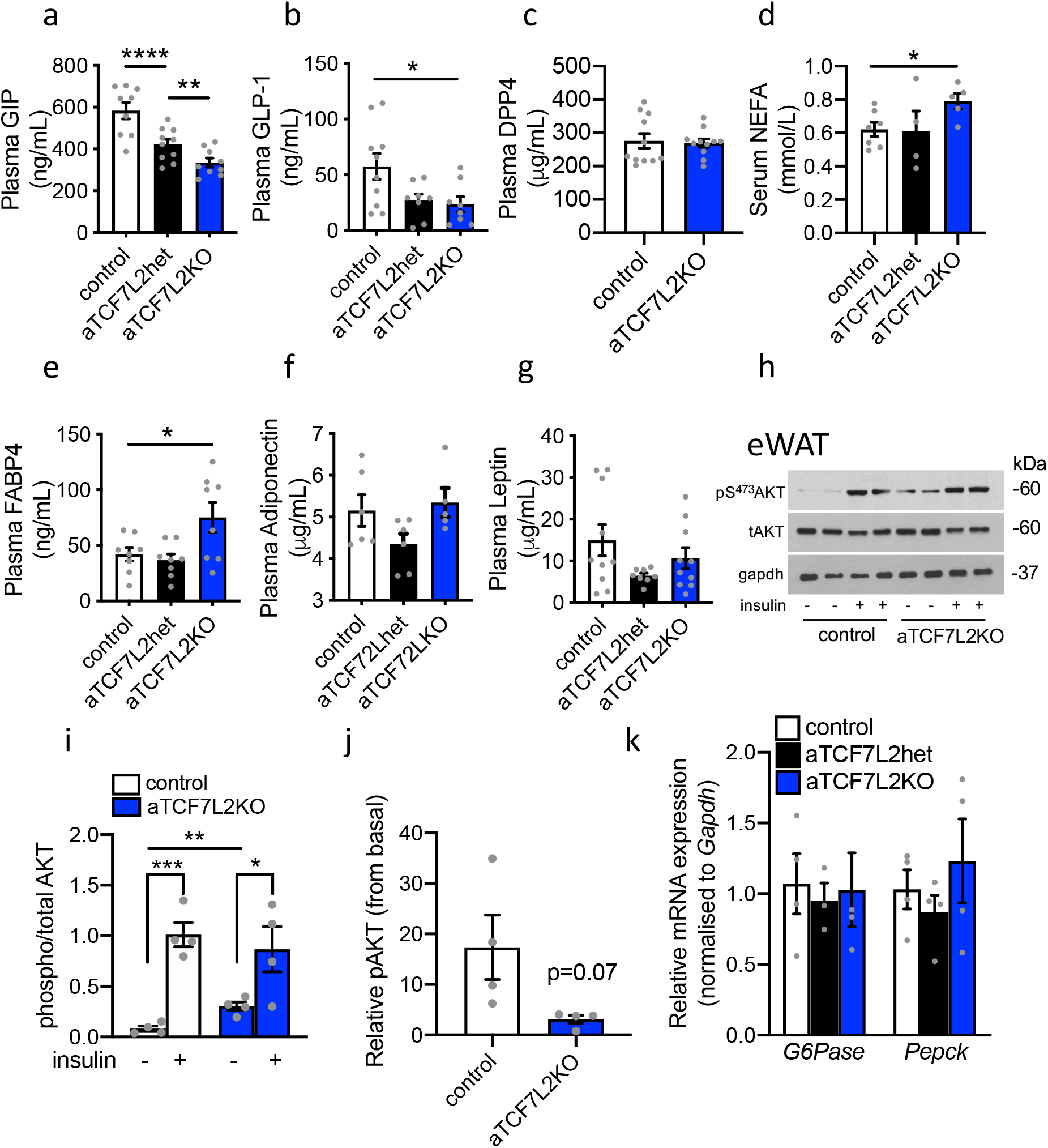
Plasma fatty acid and incretins levels depend on adipocyte *Tcf7l2* expression. (**a**) GIP plasma circulating levels. One-way ANOVA Tukey post-hoc test ****p<0.0001, **p<0.01 v control. (**b**) Total GLP-1 plasma circulating levels. One-way ANOVA followed by Tukey post-hoc test *p<0.05 versus control. (**c**) DPP4 plasma circulating levels. (**d**) NEFA serum circulating levels. One-way ANOVA followed by Tukey post-hoc test *p<0.05 versus control. (**e**) FABP4 plasma circulating levels. One-way ANOVA followed by Tukey post-hoc test *p<0.05 versus control. (**f**) Adiponectin plasma circulating levels. (**g**) Leptin plasma circulating levels. (**h**) Representative western blot of phosphorylated and total AKT from eWAT homogenates harvest from male mice 10 minutes after intraperitoneal injection of sterile PBS (-insulin) or 1 UI/kg of insulin (+ insulin) for 4 mice/genotype. (**i**) Densitometry analysis of 2 independent western blot experiments with n=4 mice/genotype. Student’s t test *p<0.05 insulin group versus no insulin group in aTCF7L2KO mice, **p<0.01 no insulin group aTCF7L2KO mice versus no insulin group in control mice, ***p<0.001 insulin group versus no insulin group in control mice. (**j**) Fold change from basal (+PBS) of phosphorylated protein expression after insulin stimulation. Student’s t-test P=0.07. (**k**) mRNA expression of hepatic insulin resistance markers in whole liver. Each dot represents one mouse. Data shown as mean ± SEM.

Evidence suggests a role of Wnt/TCF7L2 signalling in the control of lipid metabolism [24, 35]. We, therefore, sought to determine whether TCF7L2 could play a role as a regulator of fatty acid release from mature adipocyte. Plasma levels of circulating NEFA and the lipid carrier FABP4 were found to be increased respectively in aTCF7L2KO mice compared to age-and sex-matched littermate controls (NEFA: 0.79 ± 0.04 vs 0.62 ± 0.04 mmol/L, respectively; FABP4: 75.0 ± 13.5 ng/mL vs 42.1 ± 6.0 ng/mL, respectively; Fig.3d and e). Finally, we investigated endocrine adipocyte function by measuring adipokines which are usually found to be affected in insulin resistance and metabolic diseases. Plasma levels of adiponectin, leptin, resistin and PAI-1 were found to be unchanged in male mice of all genotypes (Fig.4f, g and Supplemental Fig.S2a and b).

We next explored whether loss of *Tcf7l2* expression may affect insulin signalling in adipocytes, since previous reports [24, 29] described hepatic insulin resistance in a mouse model of ablation of TCF7L2. PKB/Akt Ser473 phosphorylation was elevated at basal condition prior insulin stimulation in older (17 weeks) aTCF7L2 male mice (Fig.4h and i), but PKB/Akt Ser473 phosphorylation in response to insulin was similar adipocytes from aTCF7L2KO mice and control littermates. However, relative expression of phosphorylated Akt after insulin stimulation compared to basal appeared decreased in aTCF7L2KO mice compared to controls but was not robust enough to reach statistical significance (p=0.07; Fig.4j). Indicative of unaltered hepatic insulin sensitivity, liver levels of the gluconeogenic genes *G6Pase* and *Pepck (Pck1)* did not differ between control and aTCF7L2KO mice (Fig.4j). Therefore, our results suggest that TCF7L2 could control lipolysis and lipid metabolism in adipocyte.

### Effects of high fat diet on adipocyte-selective deletion of Tcf7l2

In order to investigate whether nutritional status had an impact on glucose metabolism in the absence of *TCF7L2* expression in adipocytes, we maintained aTCF7L2KO male mice on high fat (60%) diet for 12 weeks. We focused on males as no change were observed in female mice on chow diet. Body weight trajectory showed a similar increase in aTCF7L2KO compared to control mice (Fig.5a). Glucose tolerance was altered after intraperitoneal injection of a high concentration (2g/kg) of glucose by a delay of 15 minutes in the blood glucose peak after injection of glucose compared to controls (17.6 ± 1.5 vs 23.7 ± 2.2 mmol/L respectively; Fig.5b and c). No significant change in oral glucose tolerance (Fig.5d and e), or insulin sensitivity (Fig.5f), was observed between mice of all genotypes. Insulin secretion was impaired during i*n vivo* oral glucose challenge in aTCF7L2KO compared to littermate controls (at 15 minutes, 4.6 ± 0.2 vs 8.9 ± 1.0 ng/mL respectively; Fig.5h). We observed no robust changes following intraperitoneal injection of glucose to reach statistical significance (Fig.5g). *Ex vivo* insulin release in response to high glucose (17 mM), GLP-1 (20nM) and KCl (30 mM) was found to be no different between islets of Langerhans isolated from aTCF7L2KO mice and littermate control mice (Fig.5i).

**Fig. 5.**
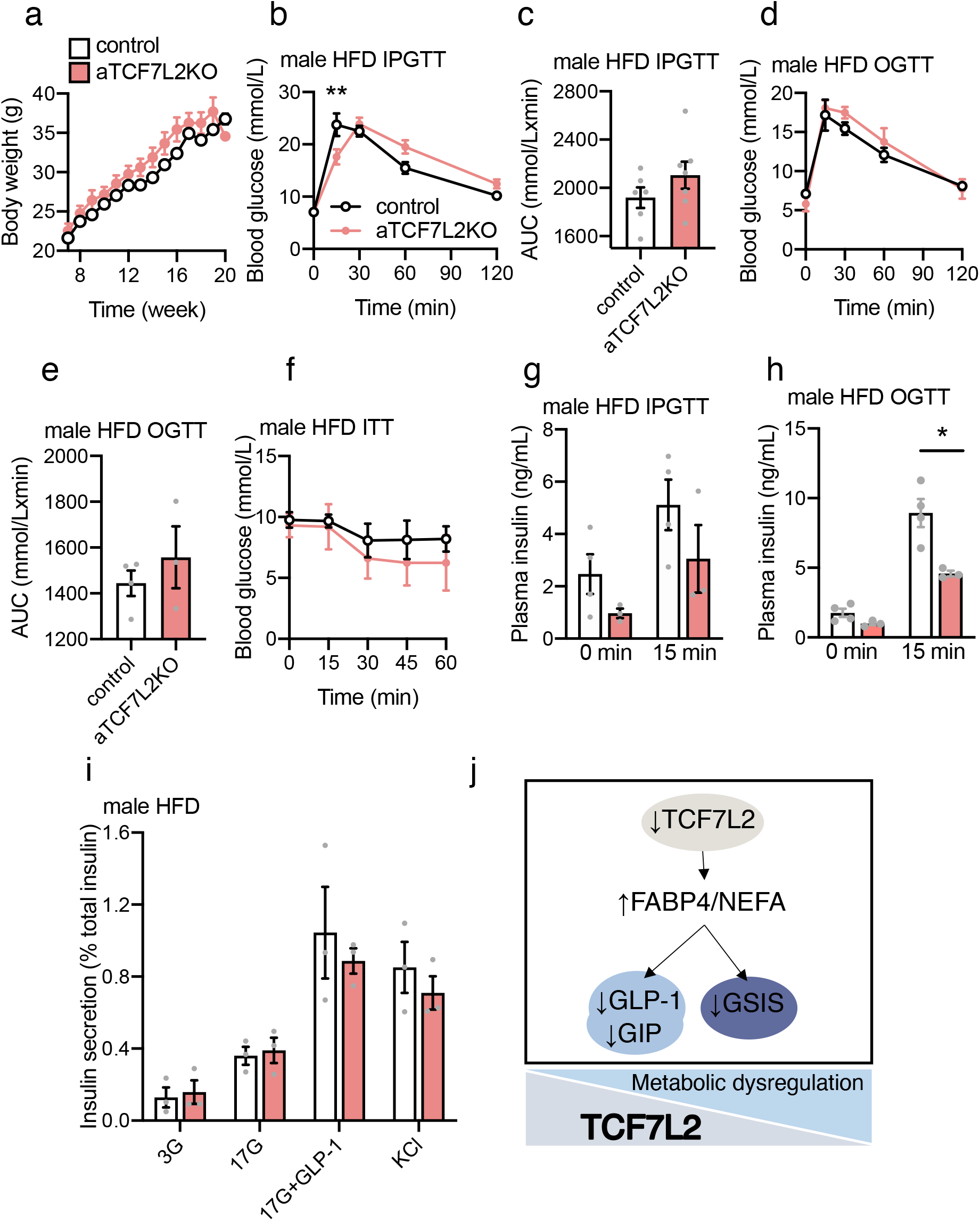
Effects of high fat diet on adipocyte-selective deletion of *Tcf7l2*. (**a**) Body weight in males during high fat diet (HFD) feeding (n=6 mice/genotype, except at 20 weeks: n=4 control mice, n=3 aTCF7L2KO mice). (**b**) Blood glucose levels during IPGTT in male mice after 9 weeks of HFD (n=6 mice/genotype). Data were analysed by two-way ANOVA followed by Bonferroni post-hoc test *p<0.05 versus control. (**c**) Area under the curve (AUC) corresponding to the IPGTT. (**d, e**) Blood glucose levels during OGTT in male mice after 12 weeks of HFD (n=4 control mice, n=3 aTCF7L2KO mice). (**f**) Blood glucose levels during IPITT in male mice after 12 weeks of HFD (n=3 mice/genotype). (**g**) Insulin plasma levels after intraperitoneal injection of glucose (2g/kg) in male mice following 12 weeks of HFD (n=4 control mice, n=3 aTCF7L2KO mice). (**h**) Insulin plasma levels after oral administration of glucose (2g/kg) in male mice following 12 weeks of HFD (n=4 control mice, n=3 aTCF7L2KO mice). Data were analysed by unpaired Student’s t-test, p*<0.05 versus control. (**i**) Insulin secretion on isolated islets from male mice after 12 weeks of HFD during static incubation of 3 mmol/l glucose (3G), 17 mmol/l glucose (17G), a combination of 17 mmol/l glucose and 20 nmol/l GLP-1 (17G+GLP-1) and 30 mol/l KCl, (n=3 mice/genotype). (**j**) Proposed mechanism: Decreased TCF7L2 expression in adipocyte provoked increased circulating levels of lipids and subsequently, decreased incretins and glucose-stimulated insulin secretion (GSIS) were observed *in vivo*. As TCF7L2 expression decreased in adipocyte, features of metabolic disorder were observed at the whole-body level. Data shown as mean ± SEM.

## Discussion

In the present study, we demonstrate that changes in the expression of *Tcf7l2* in murine adult adipose tissue may lead to alterations not only in adipocyte function but also in the function of other tissues involved in the regulation of energy homeostasis in a gender-, age-, and nutritional status-dependent manner. Thus, we provide evidence that deletion of *Tcf7l2* in adipocytes leads to alterations in the function of adipocytes, pancreatic islet beta cells, and enteroendocrine cells, thereby highlighting a role for TCF7L2 in systemic glucose homeostasis.

Wnt signalling and its effectors beta-catenin and TCF7L2 are critical during adipogenesis [6, 8, 29]. The presence of this signalling module during adulthood suggests that it may also be important for the function of adult adipocytes. In the present study, we found that young mice presented no alteration of glucose tolerance or body composition, regardless of the gender, in the absence of TCF7L2 in adipocyte, whilst defects appear with age (Fig.1g and Fig.2a). Recently, Tian et al. revealed crosstalk between Wnt signalling and females hormones through TCF7L2 to regulate lipid metabolism [35]. Thus, gender differences observed when expressing a dominant-negative form of TCF7L2 in hepatocytes would suggest that sex hormones regulate glucose homeostasis via repressing hepatic gluconeogenesis and regulate lipid metabolism [35]. In our study, we also found that female mice were largely unaffected by the loss of TCF7L2 function in the adipocyte, suggesting a role for female hormones to maintain glucose homeostasis.

The key effector of the Wnt signalling pathway is the association of beta-catenin and a member of the TCF family which may include TCF7L2, TCF7, TCF7L1 and LEF-1 [1]. The availability of free beta-catenin entering the nucleus to bind TCF7L2 is crucial for activation of expression of downstream target genes. However, the regulation of TCF7L2 expression is also essential. High-fat feeding modulates TCF7L2 expression in pancreatic islets, in hepatocytes and in human adipocytes [27, 36, 37]. Furthermore, studies have suggested that variation in TCF7L2 expression altered glucose metabolism and induces type 2 diabetes phenotype [38]. We found that deletion of a single *Tcf7l2* allele generates distinct features of obesity-induced glucose intolerance while biallelic *Tcf7l2* deletion reveals a disruption of endocrine signalling molecules. Therefore, alterations induced by deletion of *Tcf7l2* in mature adipocytes could depend on age and on the dosage of TCF7L2 expression.

We found that under chow diet fed mice lacking TCF7L2 selectively in adipocytes displayed an impaired response to oral glucose challenge (Fig. 2d) but normal tolerance to intraperitoneal injection of the sugar (Fig. 2b). This suggests an impaired incretin effect, defined as the postprandial insulin response provoked by incretin hormones such as GLP-1 and GIP. However, insulin release in response to an oral glucose challenge was maintained (Fig. 3d), whilst glucose-stimulated insulin secretion *ex vivo* from isolated islets of Langerhans is impaired (Fig. 3e). Our data therefore suggest that a mechanism exists to maintain insulin release *in vivo* after deletion of *Tcf7l2* from adipocytes. However, when challenged with high fat feeding, impaired glucose tolerance is associated with impaired insulin secretion (Fig.5b and h). Future studies will need to assess further the effects of high-fat diet feeding on beta cell function on a larger cohort, as exploration were suspended during the pandemic events of COVID-19.

A striking finding in the present study is that TCF7L2 is required in adipose tissue for normal incretin production and insulin secretion: we reveal that decreased *Tcf7l2* expression in mature adipocytes leads to lowered circulating levels of GLP-1 and GIP (Fig.4a and b). This suggests that a compensatory effect is unmasked in aTCF7L2KO mice to stimulate insulin secretion in pancreatic beta cells when the incretin effect is compromised. One possible explanation to reconcile our findings on *in vivo* and *ex vivo* glucose-stimulated insulin secretion is that elevated fatty levels compensate in part for the lowered levels of circulating incretins, acting to amplify insulin release through the action of fatty acid receptors. NEFA and specifically long-chain fatty acids potentiate glucose-stimulated insulin secretion [39, 40]. Moreover, a direct insulinotropic action of FABP4, a cytosolic lipid chaperone expressed and secreted by white and brown adipocytes may act directly on pancreatic beta cells as demonstrated in previous studies showing that recombinant FABP4 administration enhanced glucose-stimulated insulin secretion *in vitro* and *in vivo* [41, 42]. In a small type 2 diabetes cohort, increased serum FABP4 was correlated to enhanced insulin release [43]. Taking these findings together, lowered circulating incretins may have provoked impaired glucose tolerance with no apparent change in beta cell function to release insulin *in vivo*-compensated by stimulation of glucose-stimulated insulin secretion by higher circulating FABP4 and NEFA-in aTCF7L2KO mice. Nevertheless, there is a need to further investigate the mechanisms that may link impaired glucose tolerance with normal insulin secretion. These might include mechanisms such as reduced insulin-independent glucose disposal, or insulin resistance-induced altered GLUT2 trafficking in enterocytes [44].

By what mechanisms might depletion of *Tcf7l2* from adipocytes lead to a decrease in the circulating levels of GLP-1 and GIP? Our data suggest that decreased incretin levels are unlikely to be due to an increase in the rate of degradation of these hormones in the bloodstream, as no change was found in circulating DPP4 in aTCF7L2KO mice (Fig.4c). We observed elevated circulating plasma fatty acids and FABP4 levels (Fig.4d and e), reflecting altered adipocyte metabolism in the absence of TCF7L2. Wnt and TCF7L2 are regulators of lipid metabolism in hepatocytes [35], consistent with elevated plasma triglycerides associated with *TCF7L2* risk variant rs7903146 [45]. Moreover, the genetic variant is localized near the regulatory region for Acyl-CoA Synthetase Long Chain 5 (ACSL5) which activates fatty acids to generate long chain fatty acyl CoA [45]. Consistent with our model, Martchenko and colleagues [46] have recently reported fatty acid-induced lowering of circadian release of GLP-1 from L-cells as a result of decreased *Bmal*1 expression. Similarly, Filipello et al [47] reported decreased insulin-dependent GLP-1 secretion from L-cell-derived GLUTag cells, and increased glucagon release, upon fatty acid treatment. Similar findings on GLP-1 secretion were reported by others [48, 49], whist long chain saturated (palmitate) but not unsaturated (oleate) fatty acids lead to L-cell apoptosis [50]. On the other hand, activation of free fatty acid receptors with FFAR1/GPR40 agonist TAK-875 [51] or with short-chain fatty acids (FFAR2/GPR43) [52] acutely increased GLP-1 secretion from L-cells, indicating a balance between shorter term, and more chronic “lipotoxic” effects, is likely to govern overall GLP-1 production, with the latter predominating after *Tcf7l2* deletion in adipocytes. To explore further the direct role of TCF7L2 in lipid metabolism, future studies will be necessary to assess lipolysis and insulin action in adipocyte lacking TCF7L2.

In this study, high fat diet feeding altered glucose tolerance and insulin secretion. Some discrepancies emerge from our report and the previous studies using the same genetic model. When examining the effects of conditional deletion *Tcf7l2* in mature adipocytes, Chen et al. [29] found impairments in glucose tolerance after intraperitoneal injection of glucose in 3-month-old males and females under standard diet associated with hepatic insulin resistance. Geoghegan et al. [24] also generated a conditional knockout of *Tcf7l2* in the adipocyte, reporting that knockout animals maintained on regular chow displayed no change in intraperitoneal glucose tolerance, whilst exaggerated insulin resistance and impaired glucose tolerance were apparent after high fat feeding [24]. The authors demonstrated a role for TCF7L2 in regulating lipid metabolism, finding a reduction in lipid accumulation and lipolysis during high fat feeding. We note that slightly different genetic strategies were used to create the mouse models in each case. Thus, both our own and the earlier studies used a similar same *LoxP* strategy with a *Cre* recombinase under the control of the adiponectin promoter, but we have deleted exon 1, whereas Chen et al. targeted exon 11 and Geoghegan et al. targeted exon 5 [24, 29]. The *Tcf7l2* gene consists of 17 exons [53] and tissue-specific splicing variants could exert tissue-dependent distinct function and impact differently on specific cell type function such as the adipocyte or the beta cell [54]. Alternative splicing of *Tcf7l2* potentially results in transcripts lacking exons 1 and 2 predicted to encode proteins lacking the β-catenin-binding domain [22]. Might changes in TCF7L2 expression in adipose tissue contribute to the effects of type 2 diabetes-associated variants? Whilst inspection of data in the GTEX database [55] does not reveal any genotype-driven alteration in subcutaneous or breast adipose tissue *TCF7L2* expression for rs7903146, we note that other intronic variants in this gene, such as rs6585195, reveal significant expression quantitative trait loci for *TCF7L2*, and may conceivably influence incretin and insulin levels through the mechanisms identified here.

In summary, TCF7L2 could play a post-developmental role on metabolic tissues depending on its level of expression and the age of the mice. Our findings provide an unexpected insight into the action of TCF7L2 and reveal a novel mechanism through which adipocytes may impact insulin and incretin secretion. In demonstrating an action on multiple tissues, we also reveal a new level of complexity in the effects of gene action at the systems level on disease risk. Such findings may highlight the importance of early interventions with incretins when TCF7L2 expression is compromised and therefore the development of novel personalized therapies.

## Abbreviations

DPP4: dipeptidyl peptidase 4
eWAT: epididymal white adipose tissue
FABP4: fatty acid binding protein 4
GIP: glucose-dependent insulinotropic polypeptide
GLP-1: Glucagon-like peptide-1
GSIS: glucose-stimulated insulin secretion
HFD: high fat diet
HTRF: Homogeneous Time Resolved Fluorescence
iWAT: inguinal subcutaneous white adipose tissue
KO: knockout
PAI-1: plasminogen activator inhibitor-1
TCF7L2: transcription factor 7-like 2.

## Acknowledgements

The authors would like to thank Dominic Withers (MRC London Institute of Medical Sciences (LMS) at Imperial College London) for the Adipoq-Cre mouse, Lorraine Lawrence (Research Histology Facility at Imperial College London) and Stephen Rothery (Facility for Imaging by Light Microscopy FILM at Imperial College London) for technical assistance.

## Contribution statement

M-SN-T co-designed the study, collected, analysed, interpreted the data and drafted the manuscript. GdSX conceived and co-designed the study, collected, interpreted the data and substantially critically revised the manuscript. GAR conceived the study and co-wrote the manuscript. AM-S contributed to the collection of data and critically revised the manuscript for important intellectual content. IL contributed for resources. All authors gave final approval of the manuscript and gave consent to publication. GAR is the guarantor of this work.

## References

[1] Jin T, Liu L (2008) The Wnt signaling pathway effector TCF7L2 and type 2 diabetes mellitus. Mol Endocrinol 22(11): 2383–2392. 10.1210/me.2008-0135

[2] da Silva Xavier G, Mondragon A, Sun G, et al. (2012) Abnormal glucose tolerance and insulin secretion in pancreas-specific Tcf7l2-null mice. Diabetologia 55(10): 2667–2676. 10.1007/s00125-012-2600-7

[3] Mitchell RK, Mondragon A, Chen L, et al. (2015) Selective disruption of Tcf7l2 in the pancreatic beta cell impairs secretory function and lowers beta cell mass. Hum Mol Genet 24(5): 1390–1399. 10.1093/hmg/ddu553

[4] Boj SF, van Es JH, Huch M, et al. (2012) Diabetes risk gene and Wnt effector Tcf7l2/TCF4 controls hepatic response to perinatal and adult metabolic demand. Cell 151(7): 1595–1607. 10.1016/j.cell.2012.10.053

[5] Oh KJ, Park J, Kim SS, Oh H, Choi CS, Koo SH (2012) TCF7L2 modulates glucose homeostasis by regulating CREB- and FoxO1-dependent transcriptional pathway in the liver. PLoS Genet 8(9): e1002986. 10.1371/journal.pgen.1002986

[6] Ross SE, Hemati N, Longo KA, et al. (2000) Inhibition of adipogenesis by Wnt signaling. Science 289(5481): 950–953. 10.1126/science.289.5481.950

[7] Longo KA, Wright WS, Kang S, et al. (2004) Wnt10b inhibits development of white and brown adipose tissues. J Biol Chem 279(34): 35503–35509. 10.1074/jbc.M402937200

[8] Kennell JA, MacDougald OA (2005) Wnt signaling inhibits adipogenesis through beta-catenin-dependent and -independent mechanisms. J Biol Chem 280(25): 24004–24010. 10.1074/jbc.M501080200

[9] Palsgaard J, Emanuelli B, Winnay JN, Sumara G, Karsenty G, Kahn CR (2012) Cross-talk between insulin and Wnt signaling in preadipocytes: role of Wnt co-receptor low density lipoprotein receptor-related protein-5 (LRP5). J Biol Chem 287(15): 12016–12026. 10.1074/jbc.M111.337048

[10] Yoon JC, Ng A, Kim BH, Bianco A, Xavier RJ, Elledge SJ (2010) Wnt signaling regulates mitochondrial physiology and insulin sensitivity. Genes Dev 24(14): 1507–1518. 10.1101/gad.1924910

[11] Singh R, De Aguiar RB, Naik S, et al. (2013) LRP6 enhances glucose metabolism by promoting TCF7L2-dependent insulin receptor expression and IGF receptor stabilization in humans. Cell Metab 17(2): 197–209. 10.1016/j.cmet.2013.01.009

[12] Grant SF, Thorleifsson G, Reynisdottir I, et al. (2006) Variant of transcription factor 7-like 2 (TCF7L2) gene confers risk of type 2 diabetes. Nat Genet 38(3): 320–323. 10.1038/ng1732

[13] Strawbridge RJ, Dupuis J, Prokopenko I, et al. (2011) Genome-wide association identifies nine common variants associated with fasting proinsulin levels and provides new insights into the pathophysiology of type 2 diabetes. Diabetes 60(10): 2624–2634. 10.2337/db11-0415

[14] Loos RJ, Franks PW, Francis RW, et al. (2007) TCF7L2 polymorphisms modulate proinsulin levels and beta-cell function in a British Europid population. Diabetes 56(7): 1943–1947. 10.2337/db07-0055

[15] Lyssenko V, Lupi R, Marchetti P, et al. (2007) Mechanisms by which common variants in the TCF7L2 gene increase risk of type 2 diabetes. J Clin Invest 117(8): 2155–2163. 10.1172/JCI30706

[16] Villareal DT, Robertson H, Bell GI, et al. (2010) TCF7L2 variant rs7903146 affects the risk of type 2 diabetes by modulating incretin action. Diabetes 59(2): 479–485. 10.2337/db09-1169

[17] Pilgaard K, Jensen CB, Schou JH, et al. (2009) The T allele of rs7903146 TCF7L2 is associated with impaired insulinotropic action of incretin hormones, reduced 24 h profiles of plasma insulin and glucagon, and increased hepatic glucose production in young healthy men. Diabetologia 52(7): 1298–1307. 10.1007/s00125-009-1307-x

[18] Le Bacquer O, Kerr-Conte J, Gargani S, et al. (2012) TCF7L2 rs7903146 impairs islet function and morphology in non-diabetic individuals. Diabetologia 55(10): 2677–2681. 10.1007/s00125-012-2660-8

[19] Miguel-Escalada I, Bonas-Guarch S, Cebola I, et al. (2019) Human pancreatic islet three-dimensional chromatin architecture provides insights into the genetics of type 2 diabetes. Nat Genet 51(7): 1137–1148. 10.1038/s41588-019-0457-0

[20] Elbein SC, Chu WS, Das SK, et al. (2007) Transcription factor 7-like 2 polymorphisms and type 2 diabetes, glucose homeostasis traits and gene expression in US participants of European and African descent. Diabetologia 50(8): 1621–1630. 10.1007/s00125-007-0717-x

[21] Mondal AK, Das SK, Baldini G, et al. (2010) Genotype and tissue-specific effects on alternative splicing of the transcription factor 7-like 2 gene in humans. J Clin Endocrinol Metab 95(3): 1450–1457. 10.1210/jc.2009-2064

[22] Prokunina-Olsson L, Welch C, Hansson O, et al. (2009) Tissue-specific alternative splicing of TCF7L2. Hum Mol Genet 18(20): 3795–3804. 10.1093/hmg/ddp321

[23] Cauchi S, Meyre D, Dina C, et al. (2006) Transcription factor TCF7L2 genetic study in the French population: expression in human beta-cells and adipose tissue and strong association with type 2 diabetes. Diabetes 55(10): 2903–2908. 10.2337/db06-0474

[24] Geoghegan G, Simcox J, Seldin MM, et al. (2019) Targeted deletion of Tcf7l2 in adipocytes promotes adipocyte hypertrophy and impaired glucose metabolism. Mol Metab 24: 44–63. 10.1016/j.molmet.2019.03.003

[25] Kaminska D, Kuulasmaa T, Venesmaa S, et al. (2012) Adipose tissue TCF7L2 splicing is regulated by weight loss and associates with glucose and fatty acid metabolism. Diabetes 61(11): 2807–2813. 10.2337/db12-0239

[26] Huertas-Vazquez A, Plaisier C, Weissglas-Volkov D, et al. (2008) TCF7L2 is associated with high serum triacylglycerol and differentially expressed in adipose tissue in families with familial combined hyperlipidaemia. Diabetologia 51(1): 62–69. 10.1007/s00125-007-0850-6

[27] Justesen L, Ribel-Madsen R, Gillberg L, et al. (2019) TCF7L2 Expression Is Regulated by Cell Differentiation and Overfeeding in Human Adipose Tissue. Endocr Res 44(3): 110–116. 10.1080/07435800.2019.1573827

[28] Nobrega MA (2013) TCF7L2 and glucose metabolism: time to look beyond the pancreas. Diabetes 62(3): 706–708. 10.2337/db12-1418

[29] Chen X, Ayala I, Shannon C, et al. (2018) The Diabetes Gene and Wnt Pathway Effector TCF7L2 Regulates Adipocyte Development and Function. Diabetes 67(4): 554–568. 10.2337/db17-0318

[30] McCarthy MI, Rorsman P, Gloyn AL (2013) TCF7L2 and diabetes: a tale of two tissues, and of two species. Cell Metab 17(2): 157–159. 10.1016/j.cmet.2013.01.011

[31] Eguchi J, Wang X, Yu S, et al. (2011) Transcriptional control of adipose lipid handling by IRF4. Cell Metab 13(3): 249–259. 10.1016/j.cmet.2011.02.005

[32] Ravier MA, Rutter GA (2010) Isolation and culture of mouse pancreatic islets for ex vivo imaging studies with trappable or recombinant fluorescent probes. Methods Mol Biol 633: 171–184. 10.1007/978-1-59745-019-5_12

[33] Mitchell RK, Nguyen-Tu MS, Chabosseau P, et al. (2017) The transcription factor Pax6 is required for pancreatic beta cell identity, glucose-regulated ATP synthesis, and Ca(2+) dynamics in adult mice. J Biol Chem 292(21): 8892–8906. 10.1074/jbc.M117.784629

[34] Hodson DJ, Mitchell RK, Bellomo EA, et al. (2013) Lipotoxicity disrupts incretin-regulated human beta cell connectivity. J Clin Invest 123(10): 4182–4194. 10.1172/JCI68459

[35] Tian L, Shao W, Ip W, Song Z, Badakhshi Y, Jin T (2019) The developmental Wnt signaling pathway effector beta-catenin/TCF mediates hepatic functions of the sex hormone estradiol in regulating lipid metabolism. PLoS Biol 17(10): e3000444. 10.1371/journal.pbio.3000444

[36] Columbus J, Chiang Y, Shao W, et al. (2010) Insulin treatment and high-fat diet feeding reduces the expression of three Tcf genes in rodent pancreas. J Endocrinol 207(1): 77–86. 10.1677/JOE-10-0044

[37] Ip W, Shao W, Chiang YT, Jin T (2012) The Wnt signaling pathway effector TCF7L2 is upregulated by insulin and represses hepatic gluconeogenesis. Am J Physiol Endocrinol Metab 303(9): E1166-1176. 10.1152/ajpendo.00249.2012 10.1152/ajpheart.zh4-0578-corr.2012

[38] Savic D, Ye H, Aneas I, Park SY, Bell GI, Nobrega MA (2011) Alterations in TCF7L2 expression define its role as a key regulator of glucose metabolism. Genome Res 21(9): 1417–1425. 10.1101/gr.123745.111

[39] Latour MG, Alquier T, Oseid E, et al. (2007) GPR40 is necessary but not sufficient for fatty acid stimulation of insulin secretion in vivo. Diabetes 56(4): 1087–1094. 10.2337/db06-1532

[40] Hauke S, Keutler K, Phapale P, Yushchenko DA, Schultz C (2018) Endogenous Fatty Acids Are Essential Signaling Factors of Pancreatic beta-Cells and Insulin Secretion. Diabetes 67(10): 1986–1998. 10.2337/db17-1215

[41] Wu LE, Samocha-Bonet D, Whitworth PT, et al. (2014) Identification of fatty acid binding protein 4 as an adipokine that regulates insulin secretion during obesity. Mol Metab 3(4): 465–473. 10.1016/j.molmet.2014.02.005

[42] Kralisch S, Kloting N, Ebert T, et al. (2015) Circulating adipocyte fatty acid-binding protein induces insulin resistance in mice in vivo. Obesity (Silver Spring) 23(5): 1007–1013. 10.1002/oby.21057

[43] Nakamura R, Okura T, Fujioka Y, et al. (2017) Serum fatty acid-binding protein 4 (FABP4) concentration is associated with insulin resistance in peripheral tissues, A clinical study. PLoS One 12(6): e0179737. 10.1371/journal.pone.0179737

[44] Tobin V, Le Gall M, Fioramonti X, et al. (2008) Insulin internalizes GLUT2 in the enterocytes of healthy but not insulin-resistant mice. Diabetes 57(3): 555–562. 10.2337/db07-0928

[45] Xia Q, Chesi A, Manduchi E, et al. (2016) The type 2 diabetes presumed causal variant within TCF7L2 resides in an element that controls the expression of ACSL5. Diabetologia 59(11): 2360–2368. 10.1007/s00125-016-4077-2

[46] Martchenko A, Oh RH, Wheeler SE, Gurges P, Chalmers JA, Brubaker PL (2018) Suppression of circadian secretion of glucagon-like peptide-1 by the saturated fatty acid, palmitate. Acta Physiol (Oxf) 222(4): e13007. 10.1111/apha.13007

[47] Filippello A, Urbano F, Di Mauro S, et al. (2018) Chronic Exposure to Palmitate Impairs Insulin Signaling in an Intestinal L-cell Line: A Possible Shift from GLP-1 to Glucagon Production. Int J Mol Sci 19(12). 10.3390/ijms19123791

[48] Vasu S, Moffett RC, McClenaghan NH, Flatt PR (2015) Differential molecular and cellular responses of GLP-1 secreting L-cells and pancreatic alpha cells to glucotoxicity and lipotoxicity. Exp Cell Res 336(1): 100–108. 10.1016/j.yexcr.2015.05.022

[49] Hayashi H, Yamada R, Das SS, et al. (2014) Glucagon-like peptide-1 production in the GLUTag cell line is impaired by free fatty acids via endoplasmic reticulum stress. Metabolism 63(6): 800–811. 10.1016/j.metabol.2014.02.012

[50] Goldspink DA, Lu VB, Billing LJ, et al. (2018) Mechanistic insights into the detection of free fatty and bile acids by ileal glucagon-like peptide-1 secreting cells. Mol Metab 7: 90–101. 10.1016/j.molmet.2017.11.005

[51] Christensen LW, Kuhre RE, Janus C, Svendsen B, Holst JJ (2015) Vascular, but not luminal, activation of FFAR1 (GPR40) stimulates GLP-1 secretion from isolated perfused rat small intestine. Physiol Rep 3(9). 10.14814/phy2.12551

[52] Tolhurst G, Heffron H, Lam YS, et al. (2012) Short-chain fatty acids stimulate glucagon-like peptide-1 secretion via the G-protein-coupled receptor FFAR2. Diabetes 61(2): 364–371. 10.2337/db11-1019

[53] Shiina H, Igawa M, Breault J, et al. (2003) The human T-cell factor-4 gene splicing isoforms, Wnt signal pathway, and apoptosis in renal cell carcinoma. Clin Cancer Res 9(6): 2121–2132

[54] Le Bacquer O, Shu L, Marchand M, et al. (2011) TCF7L2 splice variants have distinct effects on beta-cell turnover and function. Hum Mol Genet 20(10): 1906–1915. 10.1093/hmg/ddr072

[55] Consortium GT (2013) The Genotype-Tissue Expression (GTEx) project. Nat Genet 45(6): 580–585. 10.1038/ng.2653

