## Supplemental files for "Reduced expression of TCF7L2 in adipocyte impairs glucose tolerance associated with decreased insulin secretion, incretins levels and lipid metabolism dysregulation in male mice"

### Supplemental data

Figure S1: Uncut western blot images: (a and b) images for Figure 4h; (c and d) images for Figure 1b

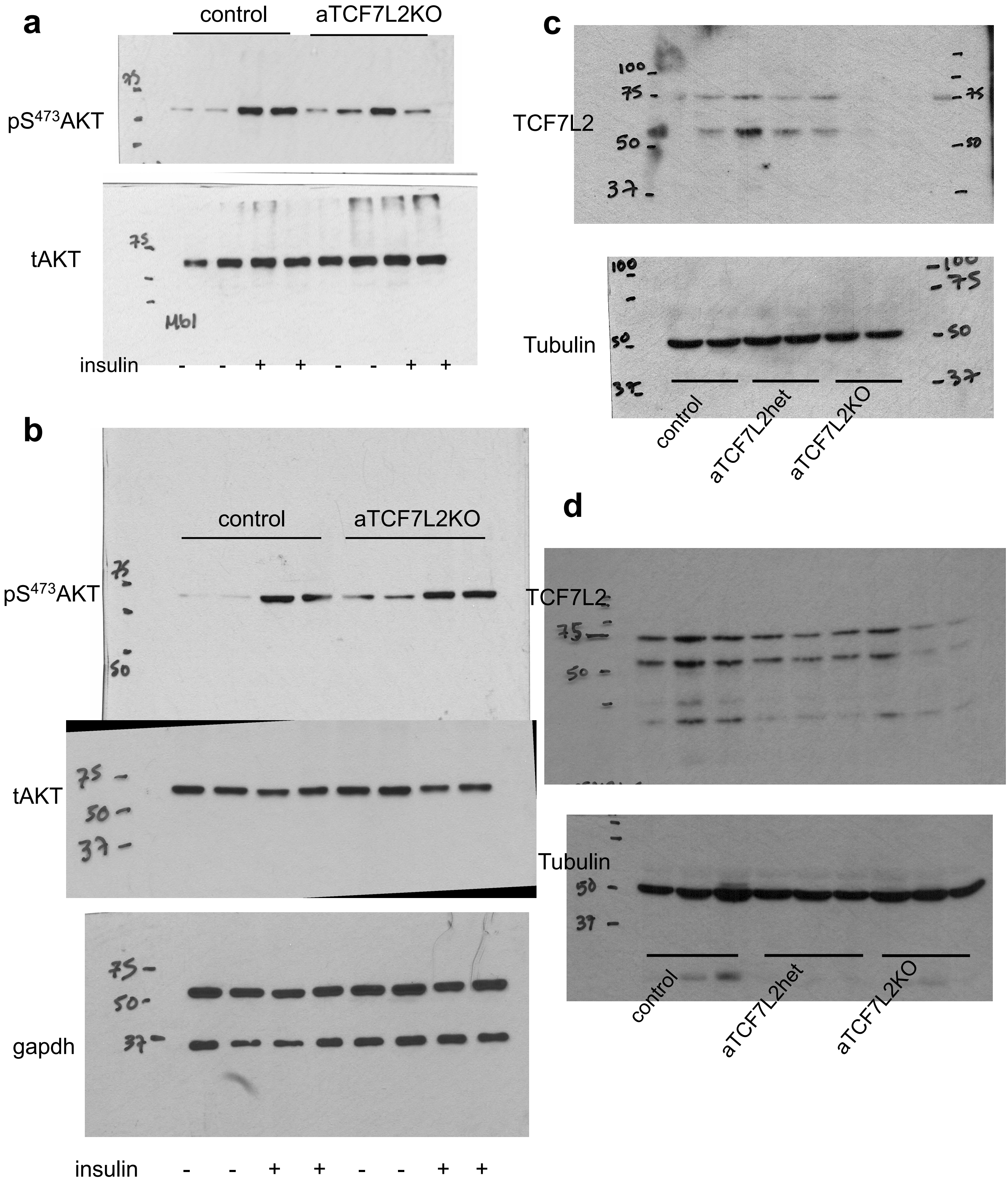

Figure S2: Plasma circulating of resistin (a) and PAI-1 (b) in male mice on chow die, Each dot represents one mouse. Data shown as mean±SEM

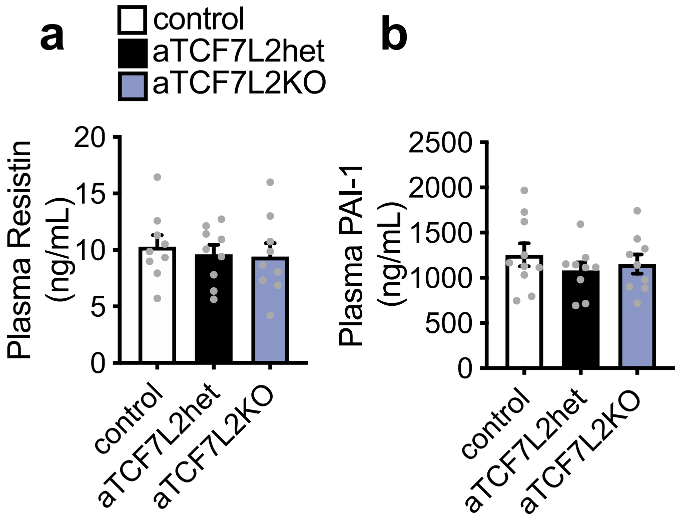


Supplemental Table 1: Primer sequences used for RT-qPCR analysis in this study.

| Target | Reverse | Forward |
|---|---|---|
| *Tcf7l2* exon 1 | CTTGGCCGCTTCTTCCAA | CAAAACAGCTCCTCCGATTCC |
| *Gcg* | CCTTCAGCATGCCTCTCAAAT | CCAAGAGGAACCGGAACAAC |
| *ins1* | AATGACCTGCTTGCTGATGGT | GCTGGTGGGCATCCAGTAA |
| *ins2* | AGCTCCAGTTGTGCCACTTGT | CGTGGCTTCTTCTACACACCC |
| *Glp1r* | ACTGCCGCCGGTATTCTCT | CCACGGTGTCCCTCTCAGA |
| *Pdx1* | CCGCCAACTTCTCGTATTTCTC | CCAAAGCTCACGCGTGGA |
| *MafA* | CCGCCAACTTCTCGTATTTCTC | CAGGTGGAGCAGCTGAAGCT |
| *G6Pase* | CGGGACAGACAGACGTTCAGC | ACTGTGGGCATCAATCTCCTC |
| *Pepck* | CTGGCTGATTCTCTGTTTCAGG | GTGCTGGAGTGGATGTTCGG |
| *Glut2/Slc2a2* | GCTTTGATCCTTCCAAGTTTGTC | TTACAGTCACACCAGCATACAC |
